# Diversity in CRISPR-based immunity protects susceptible genotypes by restricting phage spread and evolution

**DOI:** 10.1101/774349

**Authors:** Jack Common, David Walker-Sünderhauf, Stineke van Houte, Edze R Westra

## Abstract

Diversity in host resistance often associates with reduced pathogen spread. This may result from ecological and evolutionary processes, likely with feedback between them. Theory and experiments on bacteria-phage interactions have shown that genetic diversity of the bacterial adaptive immune system can limit phage evolution to overcome resistance. Using the CRISPR-Cas bacterial immune system and lytic phage, we engineered a host-pathogen system where each bacterial host genotype could be infected by only one phage genotype. With this model system, we explored how CRISPR diversity impacts the spread of phage when they can overcome a resistance allele, how immune diversity affects the evolution of the phage to increase its host range, and if there was feedback between these processes. We show that increasing CRISPR diversity benefits susceptible bacteria via a dilution effect, which limits the spread of the phage. We suggest that this ecological effect impacts the evolution of novel phage genotypes, which then feeds back into phage population dynamics.

## Introduction

Genetic diversity is a key determinant of the ecology and evolution of host-pathogen systems. Various studies of wild organisms have shown that genetic diversity within a host species often affects pathogen prevalence. The importance of diversity for disease prevalence in wild populations has been observed in numerous species, for example: cheetahs (O’Brien *et al*., 1985); Italian agile frogs (Pearman & Garner, 2005); crimson rosella parrots (Eastwood *et al*., 2017); inbred black-footed ferret populations (Thorne & Williams, 1988); inbred California sea lions (Acevedo-Whitehouse *et al*., 2003); young island populations of deer mice (Meagher, 1999); and Galapagos hawks (Whiteman *et al*., 2007). The importance of diversity for limiting disease in agricultural contexts has long been recognised (Elton, 1958), for example in rice (Zhu *et al*., 2000) and hybridising populations of honeybees (López-Uribe *et al*., 2017). In laboratory environments, more genetically diverse populations of *Daphnia magna* are more resistant to parasites (Altermatt & Ebert, 2008), an effect that depends on the genetic architecture of resistance (Luijckx *et al*., 2013). In microbial systems, *Pseudomonas aeruginosa* PA14 and *Streptococcus thermophilus* with diverse immunity alleles were shown to be more resistant against lytic bacteriophage (van Houte *et al*., 2016b; Morley *et al*., 2017). Two recent meta-analyses have shown that host diversity is a robust defence against pathogens both in agricultural (Reiss & Drinkwater, 2018) and natural populations (Ekroth *et al*., 2019).

The suggested reasons for the effect of host diversity on pathogen spread can be broadly divided into ecological and evolutionary effects. The ecological effect of diversity may manifest through a dilution effect (Ostfeld & Keesing, 2012; Civitello *et al*., 2015). The dilution effect, in terms of within-species immune diversity, is suggested to arise because increasing the number of resistant or low-quality individuals in a host population decreases the proportion of susceptible individuals in that population. This can reduce contact rates between free-living pathogens and susceptible hosts, which in turn limits the basic reproduction number of the pathogen (*R_0_*) (Dobson, 2004; Gandon, 2004; Lively, 2010). There is much observational support for the role of a dilution effect in multi-species host-pathogen systems (reviewed in (Civitello *et al*., 2015)), and some experimental work has suggested that dilution of susceptible host genotypes can limit pathogen spread in single host species systems (Dennehy *et al*., 2007; Common & Westra, 2019). Evolutionary effects of diversity may manifest through limitation of the evolutionary emergence of novel pathogen genotypes (Sasaki, 2000; Ohtsuki & Sasaki, 2006). Importantly though, the evolutionary and ecological effects of host diversity are interdependent as the basic reproductive value of a pathogen will influence its ability to evolve to overcome host resistance (Antia *et al*., 2003). Because increasing host diversity dilutes susceptible hosts, pathogens may reach smaller population sizes and generate fewer novel variants on which selection can act (Antia *et al*., 2003; Dennehy *et al*., 2006; Yates *et al*., 2006). Simultaneously, increasing host diversity can increase selection for novel variants. These two effects can maximise evolutionary emergence at intermediate host diversity (Benmayor *et al*., 2009; Chabas *et al*., 2018b).

The interaction between lytic bacteriophage (phage) and the bacterial CRISPR-Cas (Clustered Regularly Interspaced Short Palindromic Repeats; CRISPR-associated) immune system is a tractable model system to study the evolutionary ecology of infectious diseases, including the role of host diversity (van Houte *et al*., 2016b; Westra *et al*., 2017; Chabas *et al*., 2018b). CRISPR-Cas immune systems can incorporate short DNA fragments (spacers) of about 30 base pairs derived from the phage genome into CRISPR loci on the host genome (Horvath *et al*., 2008). Processed CRISPR transcripts guide Cas immune complexes to identify and cleave the invading phage genome, preventing successful re-infections (Brouns *et al*., 2008; Marraffini & Sontheimer, 2008; Garneau *et al*., 2010; Datsenko *et al*., 2012; Westra *et al*., 2012). In turn, phage can evolve to overcome CRISPR immunity by point mutation in the sequence targeted by the spacer (protospacer) or in the protospacer-adjacent motif (PAM), which flanks the protospacer and functions in self/non-self discrimination (Deveau *et al*., 2008; Mojica *et al*., 2009; Semenova *et al*., 2011; Westra *et al*., 2013). Phage evolution to overcome CRISPR immunity can lead to CRISPR-phage coevolution (Paez-Espino *et al*., 2013; Paez-Espino *et al*., 2015; Sun *et al*., 2016; Common *et al*., 2019). CRISPR loci in both natural and experimental populations can be highly diverse (Andersson & Banfield, 2008; Paez-Espino *et al*., 2013; Westra *et al*., 2015; Common *et al*., 2019), due to different bacteria in the population acquiring different spacers (Westra *et al*., 2017). Diversity has important implications for the coevolutionary interaction, as CRISPR diversity can provide increased resistance by limiting the ability of phage to acquire the mutations needed to overcome CRISPR immunity (van Houte *et al*., 2016b; Morley *et al*., 2017; Common *et al*., 2019). Phage mutation is limited by mutation supply (Lenski & Levin, 1985; Levin *et al*., 2013; Chabas *et al*., 2018a), and full phage infectivity requires mutations in all the protospacers targeted by the host CRISPR array. This becomes more difficult as individual- and population-level CRISPR diversity increases (Levin *et al*., 2013; van Houte *et al*., 2016a), and can drive rapid phage extinction (van Houte *et al*., 2016b; Morley *et al*., 2017; Chabas *et al*., 2018b).

Apart from this evolutionary effect, theory predicts that even if a phage mutant evolved that can overcome one CRISPR resistance allele in the population, its ability to amplify will be reduced in a more diverse host population (Lively, 2010). The resulting smaller phage population sizes are in turn predicted to reduce the ability of the phage to evolve to overcome other CRISPR resistance alleles in the population (Antia *et al*., 2003; Chabas *et al*., 2018b). However, these predictions remain untested. We therefore set out to explicitly test how host diversity could limit pathogen population size through a dilution effect, if and how this affects phage evolution, and the degree of interdependence between these processes. Using the bacteria *Pseudomonas aeruginosa* and its lytic phage DMS3vir, we performed an experiment where we manipulated the degree of CRISPR diversity in the host population by mixing bacterial genotypes that each carried a different CRISPR spacer. We then infected these host populations with a phage that was pre-evolved to infect only one host genotype. We then tracked changes in host fitness, and phage population dynamics and evolution, over 3 days.

## Results

To explore how population-level immune diversity would influence the population dynamics and evolution of an infective phage and its susceptible host genotype, we first developed a library of 24 *P. aeruginosa* PA14 bacteriophage-insensitive mutants (BIMs), and a corresponding library of 24 DMS3vir mutants, each of which was pre-evolved to infect only one of the 24 host BIMs (escape phage; **Fig. 1A**). We then set up an experiment where we mixed 1, 3, 6, 12 or 24 BIMs, and inoculated each mixture with ~10^6^ of a single escape phage (**Fig. 1B**). Because this experimental design increases the proportion of resistant bacterial hosts while decreasing the proportion of the susceptible host (i.e. the one that could be infected by the escape phage) (**Figure S1**), it enables us to explicitly test how this dilution effect is related to the benefits of host CRISPR allele diversity. The susceptible clone always carried a *lacZ* reporter gene, so we could follow its population dynamics and competitive performance during the experiment (**Fig. 1B**). *P. aeruginosa* Δ*pilA*, an isogenic mutant which does not have CRISPR immunity and resists phage infection via surface receptor modification, was added to each host mixture (**Fig. 1B**). *P. aeruginosa* Δ*pilA* was resistant to all phage, and allowed us to compare the fitness of the CRISPR immune populations across the CRISPR allele diversity levels. We also included 1-and 24-clone treatments inoculated with ancestral phage to which the whole bacterial population was resistant. These treatments allowed us to examine bacteria and phage dynamics at the two extremes of host diversity we tested in host populations with no pre-existing susceptibility. We then compared those dynamics with host mixtures that included a susceptible clone (van Houte *et al*., 2016b)(**Fig. 1B**). We then monitored population dynamics and evolution of the phage, of the clones with CRISPR immunity, and the susceptible fraction of the CRISPR population over 3 days.

**Fig 1.**
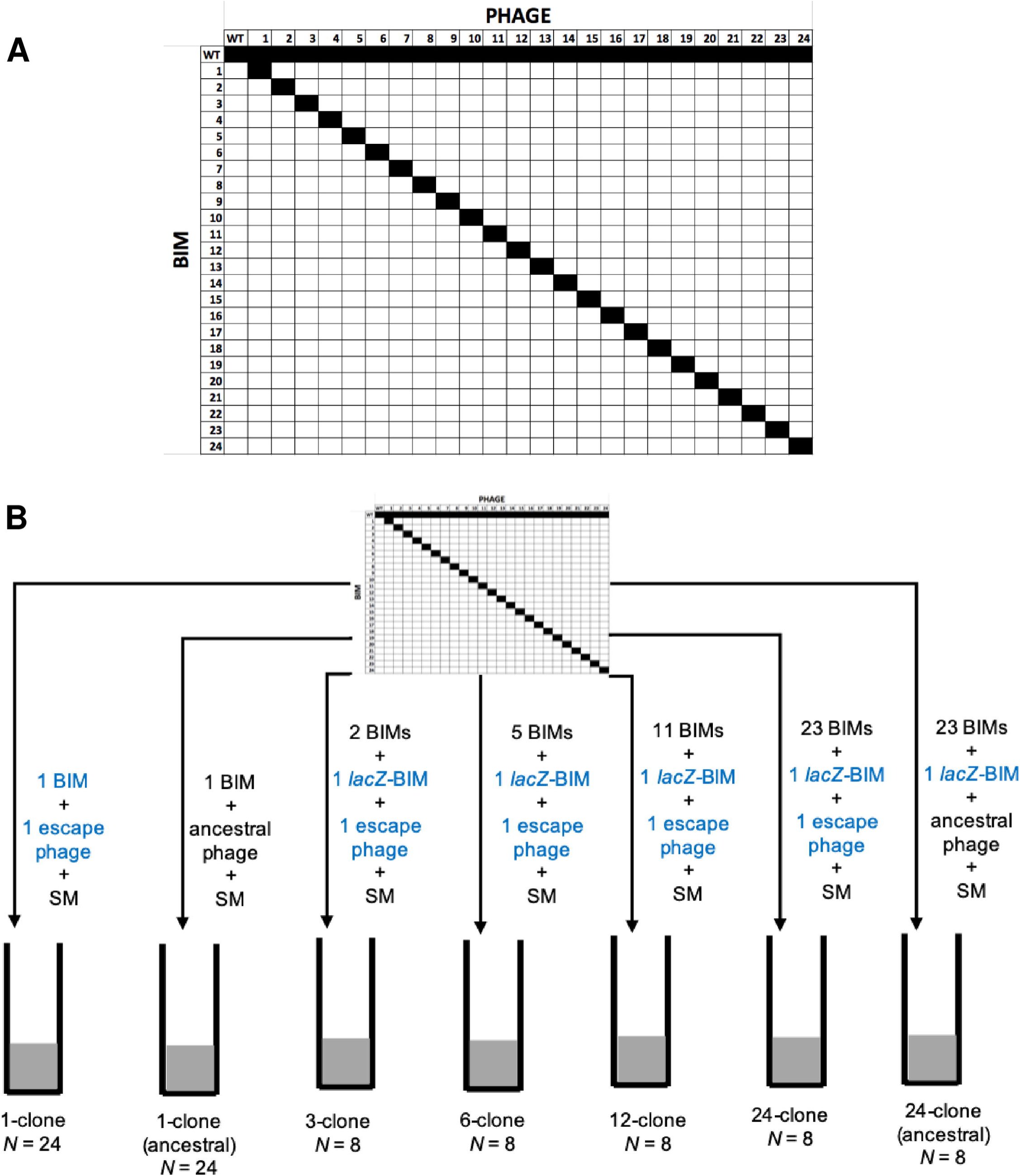
Experimental design. (A) Infectivity matrix of the library of bacteriophage-insensitive mutants (BIMs) and escape phages. The identity of BIMs and escape phage (1-24) are shown in the first row and column, respectively. Black squares represent infectivity, white indicates no infectivity. Infectivity of the wild-type DMS3vir is shown, as well as the infectivity of each escape phage on wild-type *P. aeruginosa* PA14. (B) Design of the co-culture experiment. The library of BIMs and escape phage is shown at the top. For each CRISPR allele diversity treatment, the relevant numbers of BIMs and escape phage were taken from the library and used to start the experiment, as shown by the arrows. The number of BIMs is shown next to the arrows. Blue text indicates that the BIM was engineered to carry a *lacZ* reporter gene, and that it was susceptible to the escape phage. Black text indicates that the BIMs were resistant to the phage. The number of replicates (*N*) is given underneath each treatment. SM represents the surface mutant strain PA14 Δ*pilA*.

We hypothesised that if CRISPR diversity limits phage epidemics, then phage density would be negatively correlated with CRISPR diversity. Phage densities decreased as CRISPR diversity increased (*F*_6,342_ = 48.30, *p* < 2.2 x 10^−16^; **Fig. 2**), phage population sizes decreased over the course of the experiment (regression of ln plaque-forming units [pfu] ml^−1^ over days post-infection [dpi]: *β* [95% CI] = −1.32 [-1.74, −0.94], *t*(2)_344_ = −6.40, *p* = 5.06 x 10^−10^), and phage declined faster with increasing CRISPR diversity (interaction between CRISPR diversity and days post-infection: *F*_6,314_ = 24.10, *p* < 2.2 x 10^−16^; **Fig. 2A & B**). This is consistent with an effect of CRISPR diversity that protects the bacterial population from phage. Further, phage titre in the 1-clone treatment infected with ancestral phage [“1-clone (ancestral)” in **Fig. 2B**] is statistically similar to that observed in the 1-clone treatment infected with an escape phage [“Intercept” in **Fig. 2B**], which is in line with previous data showing that monocultures composed of hosts with CRISPR immunity allow phage persistence due to rapid evolution of escape phage that overcome the CRISPR resistance allele (van Houte *et al*., 2016b). The evolution of escape phage in our monoculture treatments likely caused the observed fluctuations in the density of hosts with CRISPR immunity (**Figure S1**). When we explored the model further, we found that phage titres in the 24-clone (ancestral phage) treatment were lower than the 24-clone treatment infected with escape phage (difference in ln pfu ml^−1^: *β* [95% CI] = −2.40 [−4.67, −0.33], *t*(2)_342_ = −2.25, *p* = 0.03; **Fig. 2B**). This is consistent with a modest epidemic being able to establish by replicating on the susceptible fraction of the host population in the 24-clone treatment infected with escape phage.

**Fig 2.**
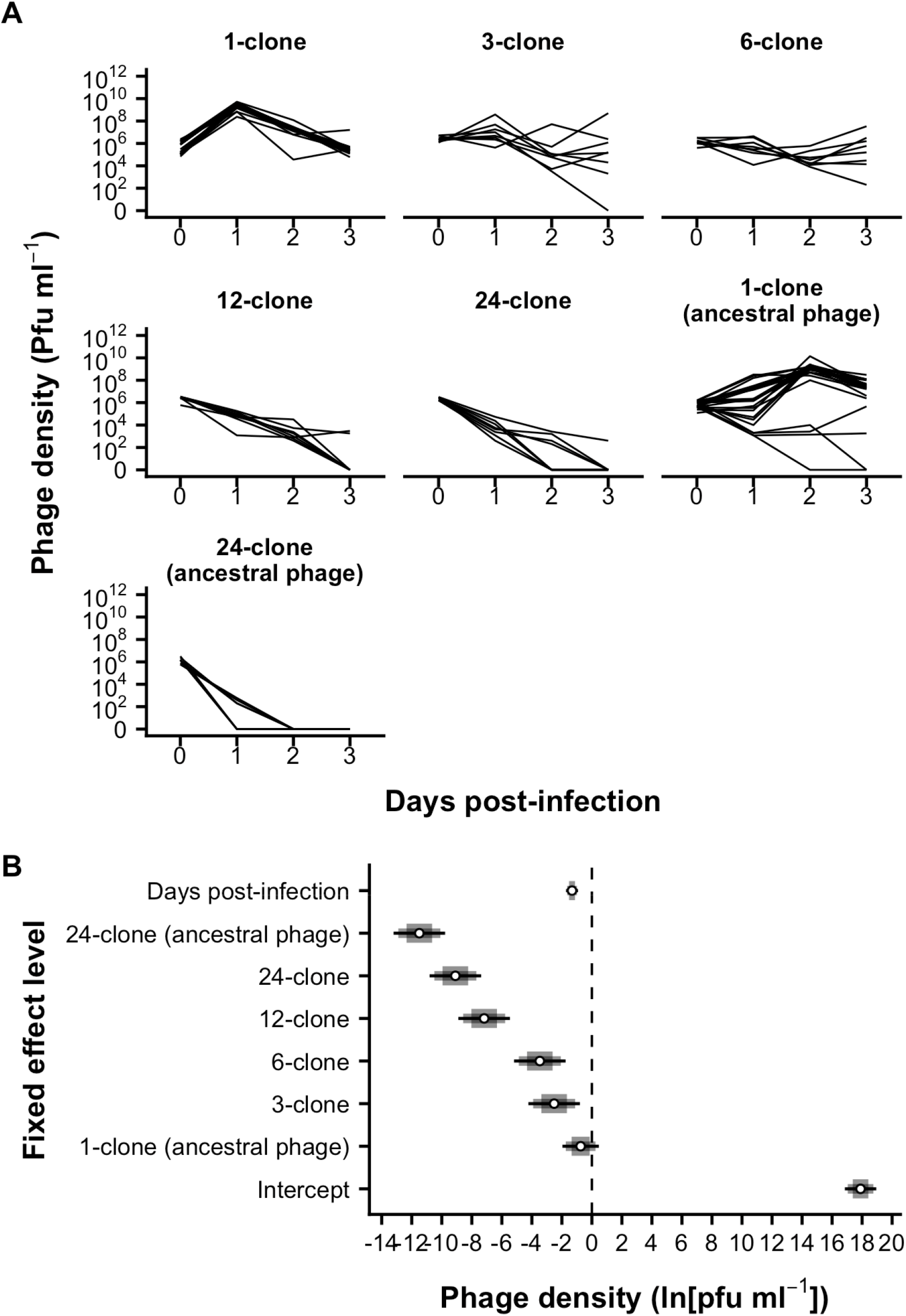
Increasing CRISPR diversity limited phage population size. (A) Population dynamics of phage at different levels of CRISPR diversity in the host population (panels). Black lines show the phage density expressed as plaque-forming units (pfu) ml^−1^ in individual replicates at each day post-infection. The limit of phage detection is 200 pfu ml^−1^. (B) Coefficients from a GLMM of the natural log of phage density (ln[pfu ml^−1^]), with CRISPR allele diversity treatment and days post-infection (dpi) as fixed effects. The intercept is the mean phage titre in the 1-clone treatment at 0 dpi. The *β* differences from the intercept are shown for the remaining levels of both fixed effects. The dotted line at zero indicates no difference from the intercept. Means are shown as white points with 67, 89% and 95% credibility intervals given in decreasing width.

Next, we hypothesised that if phage population size across our CRISPR diversity treatments was related to phage evolution, the proportion of phage that evolved an expanded infectivity range would correlate with CRISPR diversity. Given that our experiment was designed such that the proportion of the host population resistant to the escape phage increased with CRISPR diversity, there would likely have been strong selection to acquire mutations in other protospacers and PAMs to infect other hosts in the population. The evolution of range expansion did depend on diversity (Likelihood Ratio = 6.60, *p* = 0.01), and was most likely in the 6-clone treatment, particularly at 3 days post-infection (dpi) (**Fig. 3**). This result suggests that intermediate host diversity in the 6-clone treatment increased the likelihood that novel escape phage with an expanded host range would evolve (Benmayor *et al*., 2009; Chabas *et al*., 2018b). The fact that these novel phage could infect more hosts likely explains why phage population size was more consistent in the 6-clone compared to the 3-clone treatment (**Fig. 2A**), and there was no significant difference in phage population size between these treatments (**Fig. 2B**). Together, these data suggest that the increased likelihood of evolutionary emergence at intermediate host diversity balances against the potential limitation of reproduction caused by the dilution effect. These novel phage variants could then maintain phage population size.

**Fig 3.**
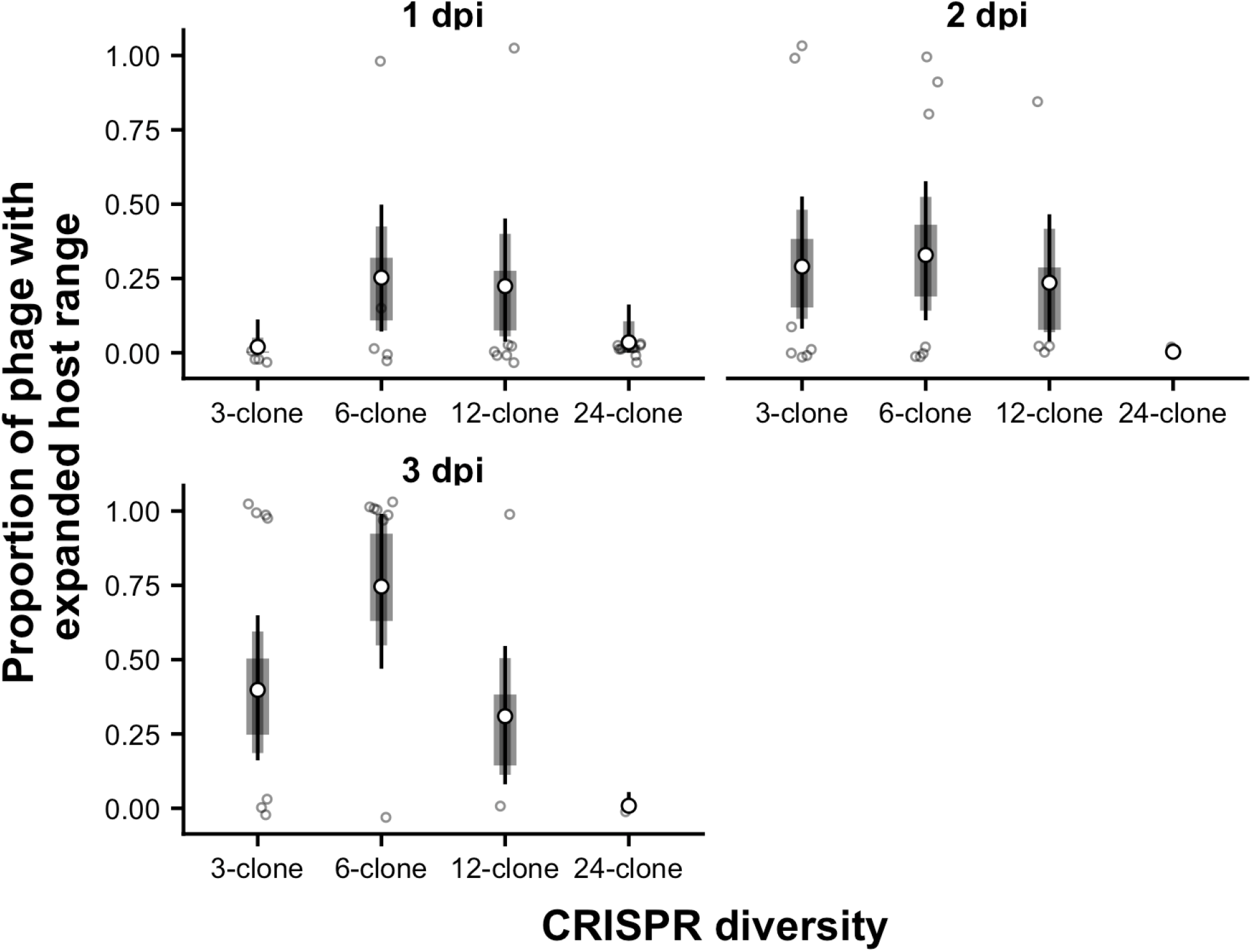
Phage evolutionary emergence was maximised at intermediate CRISPR diversity. Estimated proportion of phage that had evolved to infect a second CRISPR clone in addition to the original CRISPR clone they were pre-evolved to infect at each day post-infection in each polyclonal CRISPR diversity treatment. Because these estimates are derived from a Bayesian GLMM, they are posterior modes - not means – and are shown as white points. Similarly, 67, 89 and 95% credibility intervals are highest probability density intervals, and are given in decreasing width around mode estimates. Raw values from each replicate are shown as points.

Additionally, the presence of phage in some treatments was promoted by evolution of a host shift, where infectivity on the original host is lost but phage evolved to infect a new host. This is a less likely event as it involves two mutations: back-mutation to the ancestral state at the original protospacer followed by mutation at the new protospacer, all while incurring the cost of loss of infectivity on the original host. We observed phage in two out of eight replicates of the 24-clone treatment had evolved a host shift. Sequence data confirmed that single nucleotide polymorphisms (SNPs) in the PAM that conferred infectivity to the original host had been lost, and novel SNPs or deletions had emerged in protospacers that conferred infectivity to a new host present in the population (**Table 4 & 5 in S1 Text**). Recall that phage titres in the 24-clone (ancestral phage) treatment were lower than the 24-clone treatment infected with escape phage (**Fig. 2B**). Together with the presence of phage that evolved by host shift, these data suggest that the establishment of a modest phage epidemic in this treatment may have facilitated the evolutionary emergence of novel phage.

Given that CRISPR diversity negatively affected phage population size, we expected that this would lead to enhanced fitness of all bacterial hosts with CRISPR immunity relative to hosts without CRISPR immunity (van Houte *et al*., 2016b). We therefore hypothesised that the selection rate (Lenski, 1991; Travisano & Lenski, 1996) of all CRISPR clones (both resistant and susceptible) compared to the Δ*pilA* strain (which does not use CRISPR immunity) would be positively related to CRISPR diversity. Despite some variation among replicates, we did find that CRISPR clones in all polyclonal host populations had higher selection rates compared to clonal populations during infection with escape phage (**Figs. 4A & B**). In 3, 3, and 4 replicates in the 12-, 24-, and 24-clone (ancestral phage) treatments, respectively, the density of the Δ*pilA* strain dropped to zero (**Figure S1)**, suggesting that hosts with CRISPR immunity could occasionally outcompete those without. However, the selection rate of hosts with CRISPR immunity did not differ among the polyclonal treatments (*F*_4,113_ = 1.73, *p* = 0.15; **Fig. 4C**). Selection rate also did not change notably over time (regression of selection rate over dpi: *β* [95% CI] = 0.03 [−0.04, 0.11], *t*(2)_118_ = 0.94, *p* = 0.35) even though phage densities decreased with time.

**Fig 4.**
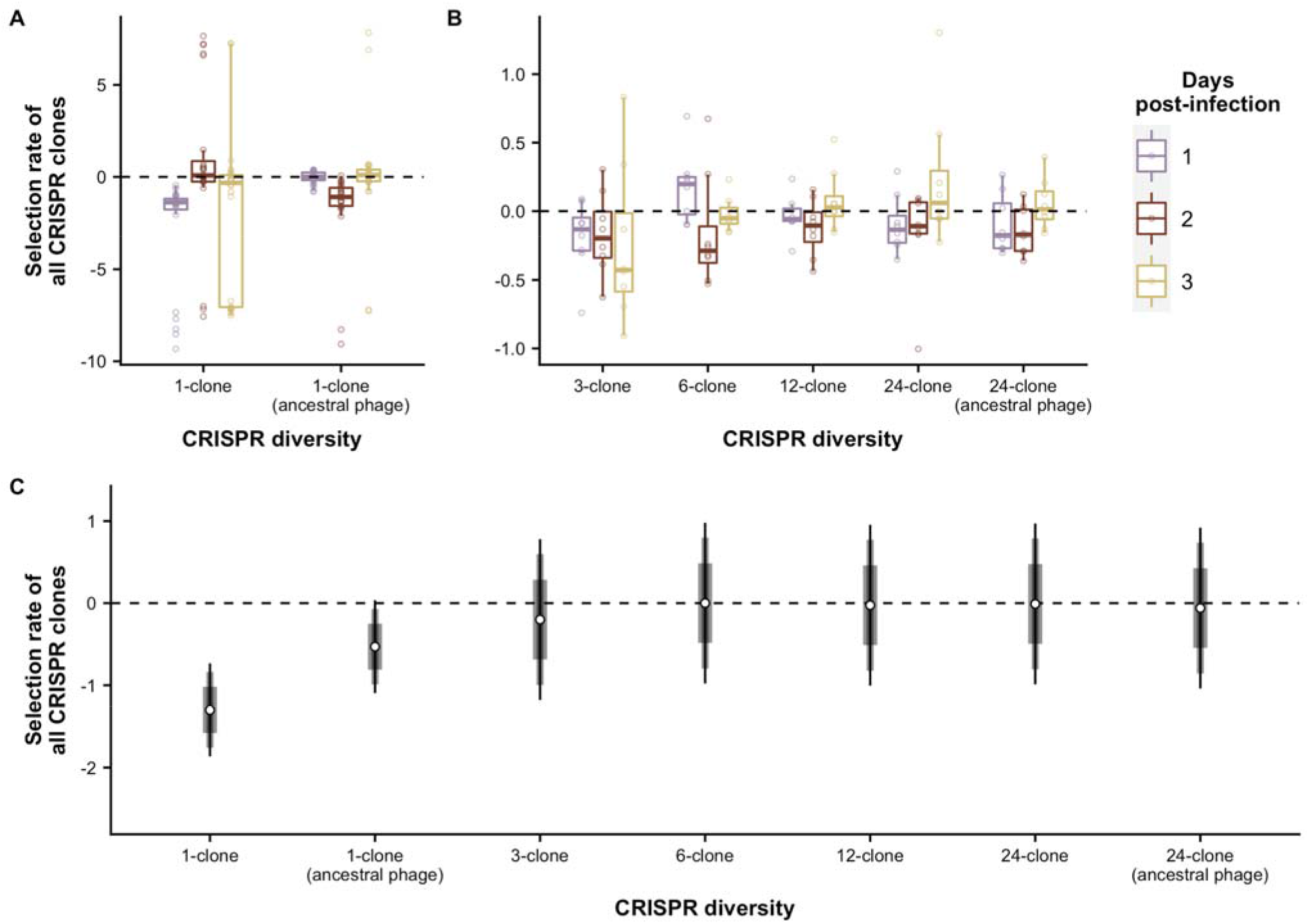
Selection rate of bacterial hosts with CRISPR immunity did not change with any level of CRISPR diversity. Selection rate of hosts with CRISPR immunity relative to a strain without CRISPR immunity in (A) the 1-clone CRISPR diversity treatments with infective and ancestral phage and (B) the remaining CRISPR diversity treatments. Days post-infection is indicated by colour. Selection rate is the natural log of the relative change in density of bacteria with CRISPR immunity (both susceptible and resistant to phage) against a strain without. The dotted line at zero indicates no difference in density change i.e. both are equally fit. Boxplots show the median, 25^th^ and 75^th^ percentile, and the interquartile range. Raw values from each replicate are shown as points. (C) Predicted mean selection rate is calculated from a GLMM that statistically controls for the effect of time. Means are shown as white points with 67, 89 and 95% credibility intervals given in decreasing width.

The absence of a detectable relationship between the level of CRISPR diversity and the selection rate hosts with CRISPR immunity relative to those without might be because escape phage dynamics were dependent on proportion of sensitive bacteria in the host population. We hypothesised that the selection rate of susceptible CRISPR clones relative to resistant CRISPR clones would be positively related to the level of CRISPR diversity, because the relative frequency of sensitive hosts declines as CRISPR diversity increases. Indeed, in some replicates in the 3- and 6-clone treatments the selection rate of the susceptible CRISPR clone was <-6, indicating that they were driven extinct by phage (**Fig. 5A**). Further, the selection rate of susceptible CRISPR clones in these treatments was on average less than zero (mean selection rate [95% CI]; 3-clone: −1.53 [−2.61, −0.45]; 6-clone: −0.99 [−2.07, − 0.09]), indicating they tended to be outcompeted by resistant CRISPR clones. Turning to the 12- and 24-clone treatments, the selection rate of susceptible clones was statistically similar to zero (12-clone: 0.12 [−0.96, 1.21]; 24-clone: 0.15 [−0.93, 1.24]; **Fig. 5B & C**), and higher than the 3- and 6-clone treatments (**Fig. 5B & C**). Together, these data show that the selection rate of susceptible CRISPR clones increased with CRISPR diversity. There was also an increase in selection rate of susceptible CRISPR clones over time among polyclonal treatments (regression of selection rate over dpi: *β* [95% CI] = 0.41 [0.03, 0.80], *t*(2)_118_ = 2.17, *p* = 0.03; **Fig. 5A & B**), which parallels the decline in phage population sizes over time in the same treatments (regression of ln pfu ml^−1^ over dpi: *β* [95% CI] = −2.58 [−3.23, −1.84], *t*(2)_114_ = −7.64, *p* = 7.27 x 10^−12^). These data suggest that susceptible hosts are protected by population-level CRISPR diversity, and that this protection is mediated by a reduced ability of phage to replicate on increasingly dilute susceptible hosts (Payne *et al*., 2018)(**Figure S1**).

**Fig 5.**
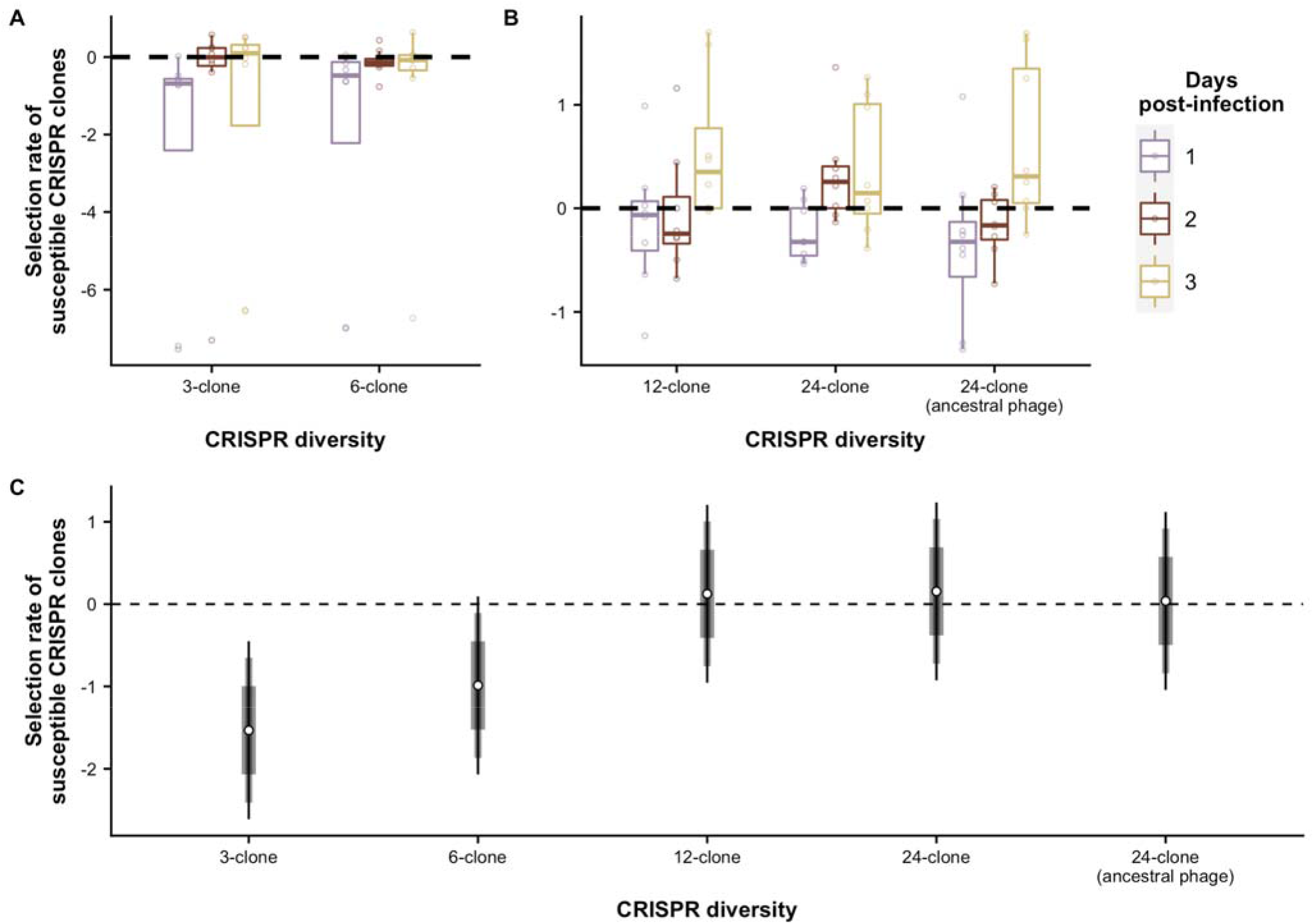
The selection rate of susceptible CRISPR clones increased with CRISPR diversity and over time. Selection rate of the susceptible CRISPR clones relative to the resistant CRISPR clones in (A) the 3- and 6-clone CRISPR diversity treatments and (B) the remaining CRISPR diversity treatments. Days post-infection is indicated by colour. Selection rate is the natural log of the relative change in density of CRISPR clones that were susceptible to phage against CRISPR clones that were resistant to phage. The dotted line at zero indicates no difference in density change i.e. both are equally fit. Boxplots show the median, 25^th^ and 75^th^ percentile, and the interquartile range. Raw values from each replicate are shown as points. The 1-clone treatments, both with infective and ancestral phage, have been excluded because all bacterial clones with CRISPR-based resistance in this treatment were susceptible, and the data is therefore present in Fig. 4A. (C) Predicted mean selection rate is calculated from a GLMM that statistically controls for the effect of time. Means are shown as white points with 67, 89 and 95% credibility intervals given in decreasing width.

## Discussion

Previous studies have shown that CRISPR diversity can limit the evolution of phage to overcome host resistance (van Houte *et al*., 2016b; Chabas *et al*., 2018b). In those studies, bacterial populations were infected with ancestral phage that had not been previously exposed to resistant hosts. Here, we examined the consequences of CRISPR diversity once a phage has already evolved to overcome one of the CRISPR resistance alleles in the population. This tractable system enabled us to closely monitor how the level of CRISPR diversity influenced the population and evolutionary dynamics of the phage, as well as the evolutionary dynamics of the host.

These analyses show that phage population sizes were larger and persisted longer in populations with lower CRISPR diversity, which is consistent with previous studies (van Houte *et al*., 2016b). Our results indicate that phage population size is increasingly limited by host diversity, because increasing host diversity reduces the proportion of susceptible individuals in the host population. This dilution effect of host resistance diversity is an important factor in explaining why genetically diverse host populations often have reduced pathogen loads (Keesing *et al*., 2006; Ostfeld & Keesing, 2012; Civitello *et al*., 2015). The finding that resistant hosts can protect the susceptible fraction of the population is also consistent with the epidemiological concept of “herd immunity” (Fine *et al*., 2011). While herd immunity has been extensively studied in eukaryotes, recent work has suggested it may be important in understanding bacteria-phage dynamics (Payne *et al*., 2018), a suggestion that for which our results provide further experimental support. This result also qualitatively matches theoretical predictions of how host diversity should affect pathogen spread in a system with matching allele genetic architecture (Lively, 2010).

In theory, this kind of ecological effect can in turn shape the evolutionary dynamics of the bacteria-phage interaction, since smaller phage population sizes will decrease the mutation supply and hence the evolutionary potential of the phage (Antia *et al*., 2003; Dennehy *et al*., 2006; Yates *et al*., 2006). We did observe that the evolution of phage host range expansion peaked at intermediate levels of CRISPR diversity. This is likely because increasing CRISPR diversity dilutes susceptible hosts, which results in smaller phage population size and hence less genetic variation on which selection can act. Simultaneously, increasing host diversity can increase selection for escape mutations. These two effects in dynamic tension can maximise evolutionary emergence at intermediate host diversity (Chabas *et al*., 2018b). The kind of interdependence between ecological and evolutionary dynamics of phage suggested by this study provides important context for other factors that can affect – and are affected by - evolutionary emergence. For example, the ability of a phage to escape a particular CRISPR spacer can influence host population structure by determining colony heterogeneity (Pyenson & Marraffini, 2020). Another example is when CRISPR immunity is incomplete, because phage with anti-CRISPR (*Acr*) genes can vary in the degree to which they inhibit the CRISPR immune response (Landsberger *et al*., 2018). Differences in the degree of CRISPR immunity because of *Acr* diversity have been shown to affect the evolutionary emergence of escape phage (Chevallereau *et al*., 2020). Understanding of how these factors interact with CRISPR allele diversity to affect the emergence and spread of escape phage will be important in future studies of CRISPR ecology and evolution.

In this study, we focussed on host populations where different CRISPR resistance genotypes were at equal starting frequencies, but natural communities are often composed of a few very common and many rare variants (Pachepsky *et al*., 2001; McGill *et al*., 2007). This likely matters for the observed dynamics, since the proportion of susceptible hosts has a large impact on the probability of evolutionary emergence of pathogens (Chabas *et al*., 2018b). Also, we focussed our analysis on the simple case where a diverse host population is infected by a clonal pathogen population. In nature, pathogen populations will frequently be genetically diverse as well (Hudson *et al*., 2006; Telfer *et al*., 2010), and increased levels of pathogen diversity may affect the benefits of host diversity (Ganz & Ebert, 2010). Indeed, previous studies of CRISPR-phage interactions suggest that infection by two different phages can increase bacteria-phage coexistence compared to infections with a single phage (Paez-Espino *et al*., 2013; Paez-Espino *et al*., 2015). Further, infection by diverse phage can select against CRISPR immunity in favour of surface-based resistance at the level of the host population, but also select for individual CRISPR clones with a more diverse array of spacers (Broniewski *et al*., 2020). The empirical system used in this study offers both a unique ability to link genotypes and phenotypes, tight experimental control over the infectivity matrix of the host-phage interaction, and a clear link with theoretical predictions. These features will make it an ideal system for more detailed studies to understand how the composition of the host population and the relative diversity levels of the phage and host shape coevolutionary interactions.

## Materials & Methods

### Bacterial strains and phage

Evolution experiments were carried out using *Pseudomonas aeruginosa* UCBPP-PA14 (which has two CRISPR loci, CRISPR1 and CRISPR2), UCBPP-PA14 Δ*pilA* (this strain lacks the pilus, which is the phage DMS3 receptor, and therefore displays surface-based instead of CRISPR-based resistance) and phage DMS3vir (Zegans *et al*., 2009). We used *P. aeruginosa* UCBPP-PA14 *csy3::lacZ* (Cady *et al*., 2012), which carries an inactive CRISPR-Cas system, for phage amplification, and for top lawns in phage spot and plaque assays. *P. aeruginosa* PA14 *csy3::lacZ, Escherichia coli* DH5α (NEB), *E. coli* CC118 λpir (NEB), and *E. coli MFDpir* (Ferrieres *et al*., 2010) were used for molecular cloning.

### Co-culture experiment

To control the levels of CRISPR diversity in our evolution experiments, we established a library of 24 *P. aeruginosa* PA14 clones each carrying a single spacer in CRISPR2 (bacteriophage-insensitive mutants; BIMs). We also independently evolved 24 phage mutants that could infect each one BIM (escape phage).

We designed 5 treatments in which we manipulated the level of CRISPR spacer diversity, based on the BIM library: monocultures (1-clone), or polycultures consisting of 3, 6, 12 and 24 clones. In order to monitor the population dynamics and relative fitness of individual bacterial clones within the polyculture treatments over the course of the experiment, we transformed 8 BIMs to carry a *lacZ* reporter gene. The *LacZ* gene encodes the β-galactosidase enzyme that hydrolyses X-gal, resulting in the production of a blue pigment. For each of the polyclonal treatments, a single BIM carrying the *lacZ* reporter gene was included. 8 BIMs were chosen for transformation so that a single clone could be monitored in each of the 3-clone mixtures (BIMs 1, 4, 7, 10, 13, 16, 19, and 22; see **Table S1**)

From fresh overnight cultures of each BIM, we made mixes of equal proportion of each clone corresponding to the diversity treatments. To monitor the population dynamics and competitive performance of all bacterial clones with an active CRISPR system, we also added PA14 Δ*pilA* (which is fully resistant to phage DMS3vir via surface-based resistance and has a distinct “smooth” colony morphology) to each mix in equal proportion to the CRISPR-carrying fraction of the population. We then inoculated 6ml of M9 minimal media (supplemented with 0.2% glucose; M9m) with bacteria from the mixed overnight cultures at a dilution of 1:100. Approximately 1×10^6^ pfu ml^−1^ of the escape phage targeting the labelled BIM were then added to each vial. We also established 1- and 24-clone treatments with ancestral phage as controls. Polyclonal treatments consisted of 8 biological replicates (*N*=8) to ensure that both BIMs and phage were equally represented across treatments, while the 1-clone treatments consisted of 24 biological replicates (*N*=24). Glass vials were incubated at 37°C while shaking at 180rpm. At 1, 2, and 3 days-post infection (dpi), the sampling of the phage and bacterial culture was repeated as described below. Cultures were transferred 1:100 to fresh media after sampling had been carried out. The experiment was terminated at 3 dpi.

Sampling proceeded as follows. Each day 180μl of culture was taken from each vial and phage was extracted using chloroform. Phage titres were determined by serially diluting extracted phage in 1x M9 salts, and then spotting 5μl of each dilution on a top lawn of *P. aeruginosa* PA14 *csy3::lacZ*, which was then incubated at 37°C for 24hrs. Phage titres were calculated from this assay. The detection limit of phage spot assays is 10^2^ pfu ml^−1^. To monitor bacterial densities, culture samples were serially diluted in 1x M9 salts, and then plated on LB agar + 40μg/ml X-gal + 0.1mM IPTG, and incubated for 48hrs at 37°C. The density of SM, CRISPR and the labelled BIM was then calculated. SM were differentiated from CRISPR clones by their “smooth” colony morphology, and the labelled BIM was identified by the blue:white screen.

We assessed the competitive performance of the all BIMs relative to the Δ*pilA* strain, and the labelled BIM relative to non-labelled BIMs by calculating selection rate (*r*) each day (*t*) from 1-3 (*n*) dpi (*r_n_* = (ln [density strain A at tn/density strain A at t_n-1_] − ln[density strain B at tn/density strain B at t_n-1_])/day) (Lenski, 1991; Travisano & Lenski, 1996), which expresses competitive performance as the natural log of the relative change in density of one competitor against another. We decided to calculate selection rate instead of relative fitness (*W*) as used in Westra *et al*. (2015), because *W* is very sensitive to starting densities and hence becomes increasingly skewed at later timepoints. Applied to our data, it consequently produced impossibly high values.

### Phage evolution

We examined phage evolution during the experiment by sampling 12 individual plaques from each replicate that had detectable levels of phage from 1 to 3 dpi, which were amplified on PA14 *csy3::lacZ* overnight in LB, at 37°C and shaking at 180rpm. Phage were extracted using chloroform, and then diluted 1000-fold. Samples of each phage were then applied on lawns of each of the 24 BIMs and PA14 *csy3::lacZ*. A successful infection was indicated by a clear lysis zone on the top lawn. Phage were classified according to whether they had expanded their infectivity range (could infect the original susceptible clone and a new clone in the BIM library). Of the phages that had undergone a host shift (lost infectivity to the original clone and could only infect a new clone), we confirmed their expanded infectivity range by sequencing the old and new protospacers on the evolved phage genome (SourceBioscience, UK). We also sequenced the relevant protospacers of the pre-evolved escape phage from the BIM-phage library and ancestral DMS3vir. Primers are given in Table S3.

### Statistical analyses

All statistical analyses were carried out in R v3.5.3 (R Core Team, 2018). Generalised linear mixed models (GLMMs) were used throughout, and replicate was treated as a random effect in all models. Days post-infection (dpi) was treated as a continuous variable and CRISPR allele diversity as a categorical variable. Model selection followed a nested approach, where full versus reduced models were compared using information criteria (Burnham & Anderson, 2003, 2004), as well as focussed comparisons of observed and predicted values. The overall statistical significance of the single and interaction effects of fixed effects (that is, treatment and dpi) was assed using ANOVA, which gives an *F*- and associated *p*-value. Simple Bayesian GLMMs from the MCMCglmm (Hadfield, 2010) package were used to analyse phage evolution to overcome model conversion issues encountered with this dataset. These models used a probit transformation (the inverse standard normal distribution of the probability) and a flat prior. The overall statistical significance of the single and additive effects of fixed effects in these models was assessed using Likelihood Ratio (LR) tests and their associated *p*-values using the VCVglmm package (Brown, 2019). Where possible, exact *p*-values are given, but R is unable to give exact values when *p* < 2.2 x 10^−16^. When phage titre was considered as the response variable, data was log-transformed to improve model fit. Credibility intervals around model coefficients and predictions were calculated to the 95%, 89% and 67% level to give the reader a clearer indication of effect size. All R code used for analyses and figures is available at this manuscript’s GitHub page.

## Acknowledgements

We thank Meaghan Castledine for help constructing the pBAM1(Gm)_lacZ plasmid.

