## Supporting Information for "Diversity in CRISPR-based immunity protects susceptible genotypes by restricting phage spread and evolution"

### Supplementary Materials & Methods

#### Library of BIMs and escape phages

11 *P. aeruginosa* PA14 bacteriophage-insensitive mutants (BIMs) that were known to have a single CRISPR2 spacer were selected from the collection of clones used in van Houte *et al.* (2016). The additional 13 BIMs were generated by evolving *P. aeruginosa* PA14 in the presence of DMS3vir. 6ml of M9 minimal media (supplemented with 0.2% glucose; M9m) was inoculated with approximately 10^6^ colony-forming units (cfu) of WT *P. aeruginosa* and 10^4^ plaque-forming units (pfu) of phage in glass vials. After 24hrs, samples from the infection were plated on LB agar. Potential CRISPR clones were identified through phenotypic and PCR analyses as described previously (Westra 2015; van Houte 2016). CRISPR amplicon sequencing (SourceBioscience, UK) confirmed that each spacer carried by a BIM was unique, so that all clones used in downstream experiments carried a different spacer. Spacer sequences were mapped against the DMS3vir genome (Genbank accession: NC_008717.1) using Geneious v9.1.8 (Kearse *et al.*, 2012) to confirm that spacers did not target overlapping regions of the phage genome. See Table S1 in Supporting Information for the spacer sequences of each BIM.

To generate 24 phage clones that could infect each BIM (escape phage), 15ml LB was inoculated with approximately 10^6^ cfu of a single BIM and approximately 10^6^ pfu DMS3vir. We also added approximately 10^6^ of *P. aeruginosa* PA14 *csy3::lacZ* to provide a pool of sensitive hosts on which phage could replicate and hence supply novel escape mutations. These cultures were then incubated overnight at 37˚C and 180rpm. Escape phage from these amplifications were identified by spot assay on a top lawn of the BIM with which they were originally mixed, and were then plaque-purified to ensure a monoclonal phage stock. Each escape phage was challenged against the entire BIM library to check for a one-to-one infection match. A successful infection was defined if a clear lysis zone was visible in the top lawn of the target BIM.

*Generating labelled BIMs*

The BIMs chosen for transformation were such that a single clone could be monitored in each of the 3-clone mixtures (that is, BIMs 1, 4, 7, 10, 13, 16, 19, and 22; see Table S1), which enabled us to measure relative frequency and fitness of a labelled BIM through time by performing a blue:white screen when plating on LB agar supplemented with 40µg/ml X-gal.

All cloning reactions to generate the labelled BIMs were carried out according to manufacturers’ instructions unless stated otherwise. Restriction enzymes, Antarctic phosphatase, and T4 DNA ligase were purchased from NEB; High-Fidelity (HF) versions were used if available. Strains, primers, and plasmids used for molecular work are outlined in Table S2. We used the synthetic mini-Tn5 transposon vector pBAMD1-6 (Martínez-García *et al.*, 2014) to deliver the *lacZ* gene to target BIMs. pBAMD1-6 is a non-replicative vector in *P. aeruginosa* encoding a Tn5 transposase, which allows for insertion of a gentamicin resistance gene (GmR) as well as any cargo genes into the bacterial chromosome. To introduce *lacZ* as a cargo gene, we amplified it from PA14 *csy3::lacZ* using primers lacZ_amp_fw and lacZ_amp_rv (Table S2) using Phusion High-Fidelity Polymerase (ThermoFisher). The PCR product was cleaned up (QIAgen PCR cleanup kit) and sub-cloned into pMA-RQ_Cas (Walker-Sünderhauf, unpublished) using NcoI-HF and KpnI-HF to generate a construct in which *lacZ* gene expression is driven by a constitutive β-lactamase promoter P3 (Genbank accession: J01749, region 4156..4233). Using standard molecular cloning protocols and restriction enzymes HindIII-HF and KpnI-HF, this promoter and the downstream *lacZ* gene was inserted into pBAMD1-6 to generate pBAM1(Gm)_lacZ. pBAM1(Gm)_lacZ was transferred into *E. coli* MFD*pir* by electroporation.

Tn5 insertions of the recipient BIMs were carried out by conjugative pBAM1(Gm)_lacZ delivery. *E. coli MFD*pir + pBAM1(Gm)_lacZ was used as donor and grown overnight in 5ml LB + 0.3mM diaminopimelic acid (DAP) + 30 µg/ml gentamicin at 37˚C, 180 rpm. Recipient BIMs were grown overnight in 5ml LB at 37˚C, 180rpm. 10ml of fresh media was inoculated from these overnight cultures, and grown at 37˚C and 250rpm until OD_600_ ~ 0.6, then pelleted and washed twice in 1x M9 salts, and resuspended in 1ml 1 x M9 salts. 600µl of donors were mixed with 200µl recipients, pelleted, and resuspended to a volume of 100µl. The entire donor-recipient mixture was pipetted onto sterile 0.2µm microfiber glass filters (Whatman) on LB agar + 0.3mM DAP plates and incubated for 2 days at 37˚C. To recover cells, filters were placed into 2.5ml LB and vortexed. 100µl of recovered cells were plated onto LB agar + 30 µg/ml gentamicin + 40µg/ml X-gal + 0.1mM IPTG plates and incubated at 37˚C for 2 days to select for BIMs with Tn5 insertions in their genome (absence of DAP selects against the donor strain).

Because Tn5 inserts at random positions in the *P. aeruginosa* genome, this may affect fitness. We therefore sampled three blue colonies of each transformed BIM and conducted 24hr competition experiments against their untransformed counterpart to verify their fitness was unaffected. The relative fitness of the transformed BIM was calculated as described previously (*W*_n_ = [(fraction transformant at t_n_) * (1 – (fraction stain transformant at t_0_)) ] / [(fraction transformant at t_0_) * (1 – (fraction transformant at t_n_)])(Westra *et al.*, 2015). If Tn5 insertion disrupted the CRISPR-Cas system, the transformed BIM would regain susceptibility to ancestral DMS3vir. We therefore checked for this by spotting ancestral DMS3vir on a top lawn of the transformed BIM. If no clear lysis zone was visible on the top lawn, we determined that the CRISPR-Cas system was functional. These checks confirmed that transformation with pBAM1(Gm)_lacZ did not affect relative fitness of the BIMs compared to their untransformed counterparts, or disrupt the CRISPR-Cas system.

#### **Supplementary Figures**

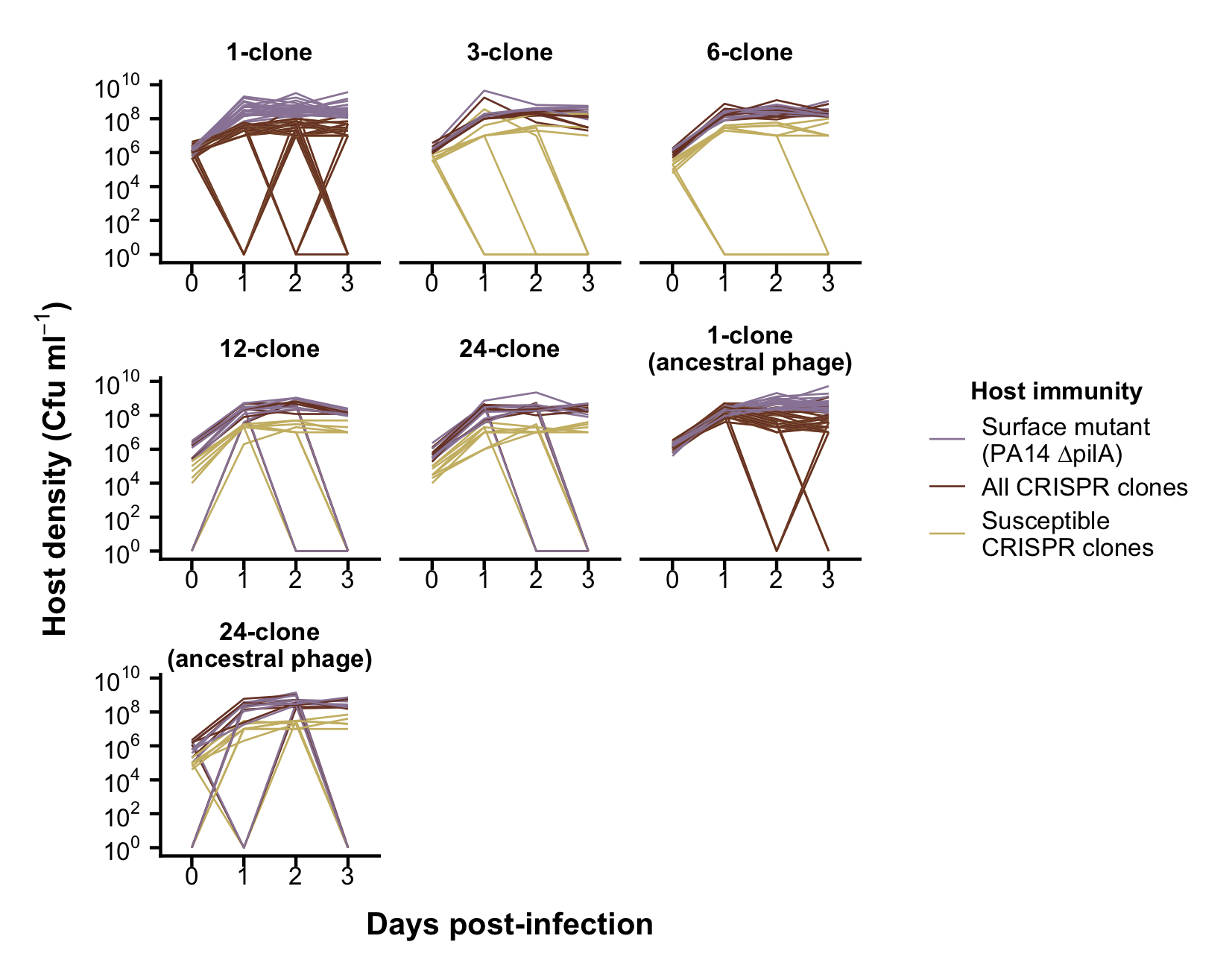

**Figure S1. Density of surface mutants, all CRISPR clones, and the susceptible CRISPR clone**

Population dynamics of the bacterial host population at different levels of CRISPR allele diversity in the host population (panels). Lines show the density expressed as colony-forming units (cfu) ml^-1^ in individual replicates at each day post-infection (x-axis). Line colours indicate host immunity: *P. aeruginosa* PA14 *∆pilA* surface mutant, which is completely resistant to phage; all CRISPR clones from the BIM library; and susceptible CRISPR clones.

### Supplementary Tables

**Table S1** Sequence of spacers in the CRISPR2 locus of each of the 24 bacteriophage-insensitive mutants (BIMs) used in the co-culture experiment. Clones transformed to carry a lacZ reporter gene using pBAM1(Gm)_lacZ are highlighted in blue.

| **BIM** | **Spacer sequence** |
| --- | --- |
| 1 | ATTTCAGTCCTTCCTGATCGCGTAGAGCCAAG |
| 2 | CATCTTCCCGCTCGATGGCGGTCAGCGTGCGC |
| 3 | CGCGTGAATGGCCCGGCGCTGAGCTGCGCTAT |
| 4 | AAGGGCATCAACCTGGCCGAAGGCGGCGCGCC |
| 5 | CGGTCGAACACGCCCTTATAGCGCTTCAGGCC |
| 6 | GATGTTCATCGCTGCCGGGCAGCGCGACATAC |
| 7 | AAACAGCGTCATGTCCAGGAGCTGCCGCTCGC |
| 8 | ACGGCAAGTTGAGTCTGGCCCTGGATGCTGAC |
| 9 | CCGGAAGTCCCGGCCGGTGTAGACGAGATAAA |
| 10 | GGCTCGACCAGGCGGCCCAGGGCGGCGTCGAT |
| 11 | TCAGGACCCCGACCAGATGGCGGCCGAAATGT |
| 12 | CGCCTGGAGACCCTGAAGGCCAATACCGAAAA |
| 13 | CCGAACGCATANANGGCGCANGGCACAGGGGT |
| 14 | ATGGGGATTCAGAGCTACGGCGATACCGCCCT |
| 15 | GAAATCGGCACCGCCACGAACCACCAGAACCT |
| 16 | GTCCAGCAGGATGCCGGCATCATCAACGAAAT |
| 17 | GGCAACGATCCCCACGAGCGGCTTTGGCACCT |
| 18 | CTCAACTCCGGCGCCGAAGACGTGATTGTCGA |
| 19 | GCGGGATCGCGGAGATAGCAGCTACGCTCGTA |
| 20 | ACTTTCACGACGACCCAGAAGCGTCGGCCGTT |
| 21 | GCGGCAGGAGCGGCAGCGGGCGGCGGCAGTT |

**Table S1 contd.**

| **BIM** | **Spacer sequence** |
| --- | --- |
| 22 | GCGATCAGNTGCGGCCAATCCGTGGACTGGGT |
| 23 | GATGGCGTCAAACTCGGCCTCCAGGCGCAGCG |
| 24 | AACCTCGCGCAGTCGTTGTCCAGCGGCATCAT |

**Table S2** Plasmids, Primers, and Strains used for molecular cloning work.

| **Plasmids** | | | |
| --- | --- | --- | --- |
| **Plasmid** | **Reference** | **Genbank accession number** | **Culture conditions** |
| pMA-RQ_Cas | Walker-Sünderhauf (unpublished) |  | 100 μg/mL Ampicillin, 50 μg/mL Gentamicin, 0.3 mM diaminopimelic acid. Needs a *pir* strain to replicate. |
| pBAMD1-6 | (Martinez-Garcia *et al.* 2014) | KM403115 |  |
| pBAM1(Gm)_lacZ | This study |  |  |
| **Primers** | | | |
| **Primer** | **Sequence (5’ 🡪 3’)** | | **Usage** |
| lacZ_amp_fw | TTACCATGGATGATTACGGAT TCACTGGCCGTCGT | | Amplification of *lacZ* from PA14 *csy3:lacZ*. Adds NcoI and KpnI restriction sites onto amplicon. |
| lacZ_amp_rv | CAGGTACCTTATTTTTGACAC CAGACCAACTGGTAATGGT | |  |

**Table S2** **contd.**

| **Bacterial Strains** | | |
| --- | --- | --- |
| **Strain** | **Reference/Supplier** | **Usage** |
| *Pseudonomas aeruginosa* PA14 *csy3::lacZ* | Zegans *et al.* 2009 | Template for *lacZ* amplification. |
| *E. coli* CC18λpir | NEB | Cloning of promoter + *lacZ* onto pBAM1(Gm) |
| *E. coli* MFD*pir* | (Ferrieres *et al.* 2010) | Donor strain for pBAM1(Gm)_lacZ delivery |
| *P. aeruginosa* PA14 BIMs | This study; Table S1 | Recipients for pBAM1(Gm)_lacZ delivery |

**Table S3** Primers used to amplify and sequence the protospacers of interest of phage that were shown to have undergone host shift (lost infectivity to the original clone and could only infect a new clone) from the phenotypic assay. The first column indicates if the protospacer was the original pre-evolved or the new protospacer, with the identity of the BIM the phage could infect shown in brackets. The primer sequence and binding direction are shown, and if the primer was used for PCR or sequencing reactions.

| **Phage** | **Sequence (5’- 3’)** | **Bind direction** | **Primer** |
| --- | --- | --- | --- |
| Original (7) | CCTGGACCTTCGCGCCGGAC | F | PCR |
|  | GAGGTGAGGTCTTCGCTTTC | R |  |
|  | GTCGCACGGAATGTTCAGCGAG | R | Sequencing |
| Original (13) | TCTGGCCAGGCGCTCACAAACAA | F | PCR |
|  | GAGCGGCTTTGGCACCTGGAAC | R |  |
|  | CCAAGTGTCGCTGCCGATCA | R | Sequencing |
| New (10) | AGCTGTCCACTGCGCTGGAC | F | PCR |
|  | CCGGAACAGATGATCCCGTT | R |  |
|  | AATGTCAGCGCGGCGGTTGC | R | Sequencing |
| New (21) | CAGCGGCATCATGGGGCTGTTTG | F | PCR |
|  | AGGTACTGAAGTTTTTGGAGGG | R |  |
|  | CCGCTGCTATCCAGACGGCC | F | Sequencing |

**Table S4** Protospacer sequences of evolved phage clones which showed host shift according to the phenotypic assay from replicate 3 of the 24-clone treatment at 1 day post-infection (dpi). The CRISPR-targeted protospacer and PAM sequences of the ancestral (WT) phage and of the pre-evolved phage are shown. Numbers 1-12 are independent phage isolates from the treatment, replicate and timepoint of interest. The second column indicates if the protospacer was the original pre-evolved or the new protospacer, with the identity of the protospacer shown in brackets. Protospacer-adjacent motif (PAM) and protospacer sequences are shown separately. SNPs and deletions, relative to WT DMS3vir, are highlighted in red.

| **24-clone, replicate 3, 1 dpi** | | | |
| --- | --- | --- | --- |
| Phage | Protospacer | PAM sequence | Protospacer sequence |
| WT DMS3vir | Original (7) | GG | CGCTCGCCGTCGAGGACCTGTACTGCGACAAA |
| Pre-evolved protospacer 7 |  | **A**G | CGCTCGCCGTCGAGGACCTGTACTGCGACAA**C** |
| 1 |  | GG | CGCTCGCCGTCGAGGACCTGTACTGCGACAAA |
| 2 |  | GG | CGCTCGCCGTCGAGGACCTGTACTGCGACAAA |
| 3 |  | GG | CGCTCGCCGTCGAGGACCTGTACTGCGACAAA |
| 4 |  | GG | CGCTCGCCGTCGAGGACCTGTACTGCGACAAA |
| 5 |  | GG | CGCTCGCCGTCGAGGACCTGTACTGCGACAAA |
| 6 |  | GG | CGCTCGCCGTCGAGGACCTGTACTGCGACAAA |
| 7 |  | GG | CGCTCGCCGTCGAGGACCTGTACTGCGACAAA |
| 8 |  | GG | CGCTCGCCGTCGAGGACCTGTACTGCGACAAA |
| 9 |  | GG | CGCTCGCCGTCGAGGACCTGTACTGCGACAAA |
| 10 |  | GG | CGCTCGCCGTCGAGGACCTGTACTGCGACAAA |
| 11 |  | GG | CGCTCGCCGTCGAGGACCTGTACTGCGACAAA |
| 12 |  | GG | CGCTCGCCGTCGAGGACCTGTACTGCGACAAA |

**Table S4 contd.**

| **24-clone, replicate 3, 1 dpi** | | | |
| --- | --- | --- | --- |
| Phage | Protospacer | PAM sequence | Protospacer sequence |
| WT DMS3vir | New (10) | GG | TAGCTGCGGCGGGACCCGGCGGACCAGCTCGG |
| Pre-evolved protospacer 7 |  | GG | TAGCTGCGGCGGGACCCGGCGGACCAGCTCGG |
| Pre-evolved protospacer 10 |  | GG | **C**AGCTGCGGCGGGACCCGGCGGACCAGCTCGG |
| 1 |  | GG | **C**AGCTGCGGCGGGACCCGGCGGACCAGCTCGG |
| 2 |  | GG | **C**AGCTGCGGCGGGACCCGGCGGACCAGCTCGG |
| 3 |  | GG | **C**AGCTGCGGCGGGACCCGGCGGACCAGCTCGG |
| 4 |  | GG | **C**AGCTGCGGCGGGACCCGGCGGACCAGCTCGG |
| 5 |  | GG | **C**AGCTGCGGCGGGACCCGGCGGACCAGCTCGG |
| 6 |  | GG | **C**AGCTGCGGCGGGACCCGGCGGACCAGCTCGG |
| 7 |  | GG | **C**AGCTGCGGCGGGACCCGGCGGACCAGCTCGG |
| 8 |  | GG | **C**AGCTGCGGCGGGACCCGGCGGACCAGCTCGG |
| 9 |  | GG | **C**AGCTGCGGCGGGACCCGGCGGACCAGCTCGG |
| 10 |  | GG | **C**AGCTGCGGCGGGACCCGGCGGACCAGCTCGG |
| 11 |  | GG | **C**AGCTGCGGCGGGACCCGGCGGACCAGCTCGG |
| 12 |  | GG | **C**ANCTGCGGCGGGACCCGGCGGACCAGCTCGG |

**Table S5** Protospacer sequences of evolved phage clones which showed host shift according to the phenotypic assay from replicate 5 of the 24-clone treatment at 2 days post-infection (dpi). The CRISPR-targeted protospacer and PAM sequences of the ancestral (WT) phage and of the pre-evolved phage are shown. Numbers 1-8 are independent phage isolates from the treatment, replicate and timepoint of interest. The second column indicates if the protospacer was the original pre-evolved or the new protospacer, with the identity of the protospacer shown in brackets. Protospacer-adjacent motif (PAM) and protospacer sequences are shown separately. SNPs and deletions, relative to WT DMS3vir, are highlighted in red.

| **24-clone, replicate 5, 2 dpi** | | | |
| --- | --- | --- | --- |
| Phage | Protospacer | PAM sequence | Protospacer sequence |
| WT DMS3vir | Original (13) | GG | TGGGGACACGGGACGCGGTAGATACGCAAGCC |
| Pre-evolved protospacer 13 |  | G**A** | TGGGGACACGGGACGCGGTAGATACGCAAGCC |
| 1 |  | GG | TGGGGACACGGGACGCGGTAGATACGCAAGCC |
| 2 |  | GG | TGGGGACACGGGACGCGGTAGATACGCAAGCC |
| 3 |  | GG | TGGGGACACGGGACGCGGTAGATACGCAAGCC |
| 4 |  | GG | TGGGGACACGGGACGCGGTAGATACGCAAGCC |
| 5 |  | GG | TGGGGACACGGGACGCGGTAGATACGCAAGCC |
| 6 |  | GG | TGGGGACACGGGACNCGGTAGATACGC**T**AGCC |
| 7 |  | GG | TGGGGACACGGGACGCGGTAGATACGCAAGCN |
| 8 |  | GG | TGGGGACACGGGACGNGGTAGATACGCAAGCC |

**Table S5 contd.**

| **24-clone, replicate 5, 2 dpi** | | | |
| --- | --- | --- | --- |
| Phage | Protospacer | PAM sequence | Protospacer sequence |
| WT DMS3vir | New (21) | GG | TGCGGCAGGAGCGGCAGCGGGCGGCGGCAGTT |
| Pre-evolved protospacer 13 |  | GG | TGCGGCAGGAGCGGCAGCGGGCGGCGGCAGTT |
| Pre-evolved protospacer 21 |  | GG | TGCGGCAG**--------------------**CGGGCGGNGGCAGTT |
| 1 |  | GG | T**--------------------**GCGGCAGCGGGCGGCGGCAGTT |
| 2 |  | GG | T**--------------------**GCGGCAGCGGGCGGCGGCAGTT |
| 3 |  | GG | T**--------------------**GCGGCAGCGGGCGGCGGCAGTT |
| 4 |  | GG | T**--------------------**GCGGCAGCGGGCGGCGGCAGTT |
| 5 |  | GG | T**--------------------**GCGGCAGCGGGCGGCGGCAGTT |
| 6 |  | GG | T**--------------------**GCGGCAGCGGGCGGCGGCAGTT |
| 7 |  | GG | T**--------------------**GCGGCAGCGGGCGGCGGCAGTT |
| 8 |  | GG | T**--------------------**GCGGCAGCGGGCGGCGGCAGTT |
